# Population Genomics of GII.4 Noroviruses Reveal Complex Diversification and New Antigenic Sites Involved in the Emergence of Pandemic Strains

**DOI:** 10.1101/668772

**Authors:** Kentaro Tohma, Cara J. Lepore, Yamei Gao, Lauren A. Ford-Siltz, Gabriel I. Parra

## Abstract

GII.4 noroviruses are a major cause of acute gastroenteritis. Their dominance has been partially explained by the continuous emergence of antigenically distinct variants. To gain insights on the mechanisms of viral emergence and population dynamics of GII.4 noroviruses, we performed large-scale genomics, structural, and mutational analyses of the viral capsid protein (VP1). GII.4 noroviruses exhibited a periodic replacement of predominant variants with accumulation of amino acid substitutions. Genomic analyses revealed (i) a large number (87%) of conserved residues; (ii) variable residues that map on the previously determined antigenic sites; and (iii) variable residues that map outside of the antigenic sites. Residues from the third pattern formed motifs on the surface of VP1, which suggested extensions of previously predicted and new uncharacterized antigenic sites. The role of two motifs (C and G) in the antigenic make-up of the GII.4 capsid protein was confirmed with monoclonal antibodies and carbohydrate blocking assays. Amino acid profiles from antigenic sites (A, C, D, E, and G) correlated with the circulation patterns of GII.4 variants, with two of them (C and G) containing residues (352, 357, 378) linked with the emergence of new GII.4 variants. Notably, the emergence of each variant was followed by a stochastic diversification with minimal changes at the antigenic sites that did not progress towards the next variant. This study provides a methodological framework for antigenic characterization of viruses, and expands our understanding of the dynamics of GII.4 noroviruses that could facilitate the design of cross-reactive vaccines.

**Importance:** Noroviruses are an important cause of viral gastroenteritis around the world. An obstacle delaying the development of norovirus vaccines is an inadequate understanding of the role of norovirus diversity in immunity. Using a population genomics approach, we identified new residues on the viral capsid protein (VP1) from GII.4 noroviruses, the predominant genotype, that appear to be involved in the emergence and antigenic topology of GII.4 variants. Careful monitoring of the substitutions in those residues involved in the diversification and emergence of new viruses could help in the early detection of future novel variants with pandemic potential. Therefore, this novel information on the antigenic diversification could facilitate GII.4 norovirus vaccine design.

## Introduction

Noroviruses are a leading cause of acute gastroenteritis affecting all ages worldwide (1). They are second only to rotavirus, although this is changing in places where rotavirus vaccination is effective (2, 3). It is estimated that norovirus is responsible for approximately 685 million infections and 200,000 deaths worldwide, with a primary public health concern in children, the elderly, and immunocompromised individuals (4–6). Norovirus outbreaks usually occur during the winter season in enclosed settings such as schools, hospitals, military facilities, and cruise ships. Because norovirus is highly contagious, outbreaks can be hard to control.

The norovirus genome is a positive-sense, single-stranded RNA molecule that is organized into three open reading frames (ORFs). ORF1 encodes for a polyprotein that is co-translationally cleaved by the viral protease into six non-structural (NS) proteins required for replication. ORF2 encodes the major capsid protein (VP1), and ORF3 the minor capsid protein (VP2). The norovirus capsid consists of 180 copies of VP1, arranged in a T=3 icosahedral symmetry. X-ray crystallography of the VP1 revealed two structural domains: the shell (S) and protruding (P). The S domain is relatively conserved and forms the core of the capsid, while the P domain is more variable and extends to the exterior of the capsid protein (7, 8). The P domain interacts with host attachment factors, namely human histo-blood group antigen (HBGA) carbohydrates, which could facilitate infection. Antibody-mediated blocking of the VP1:HBGA interaction correlates with protection against norovirus disease (9, 10). Expression of VP1 results in the self-assembly of virus-like particles (VLPs) that are structurally and antigenically similar to native virions (7, 11, 12). Given the lack of a traditional cell culture system for human norovirus, experimentally developed VLPs have been an important tool to study norovirus immune responses and vaccine design.

Norovirus is highly diverse, with at least seven genogroups (GI-GVII) and over 40 genotypes defined based on differences in their VP1 sequences (13). While over 30 different genotypes from GI, GII, and GIV can infect humans, noroviruses from the GII.4 genotype are responsible for at least 70% of infections worldwide (14). Since the mid-1990s, six major norovirus GII.4 pandemics have been recorded worldwide and were associated with the following variants: Grimsby 1995 (or US95_96), Farmington Hills 2002, Hunter 2004, Den Haag 2006b, New Orleans 2009, and Sydney 2012 (15–17). The predominance of GII.4 viruses has been linked to the chronological emergence of variants in the human population, with new variants emerging around the time that the previous declines (15). The emergence of these variants has been correlated with changes on five different variable antigenic sites (A-E) that map on the surface of the P domain; thus, new viruses can evade the human immune responses elicited to previously-circulating variants (11, 18–20). Using the recently developed cell culture system for human noroviruses (21), two of these sites have been confirmed to be involved in virus neutralization (22). Studies have shown that antibodies that map to these sites can block the interaction of VP1 with carbohydrates from the HBGA; however, the antigenic sites from several monoclonal antibodies (mAbs) with blocking activity, raised against GII.4 viruses, have not been determined (11, 12, 19, 23–25). The evolving nature of GII.4 noroviruses could challenge the development of cross-protective vaccines against noroviruses; therefore, a better understanding of the mechanisms responsible for the antigenic diversification will facilitate vaccine design.

In this study, we adopted a large-scale genomics approach to identify sites that play a role in GII.4 evolution and antigenic diversification. We (re)defined the sites involved in the antigenic make-up of GII.4 pandemic variants, and found that intra-variant diversification exhibited a stochastic pattern of evolution. Importantly, we identified three sites (amino acids 352, 357, and 378) implicated with the emergence of predominant GII.4 variants, and that could help in the early detection of the next pandemic variant.

## Results

### GII.4 Inter-variant Evolution is Characterized by the Accumulation of Substitutions in the P domain

In order to investigate the evolutionary pattern of GII.4 strains, we calculated the genetic differences of 1601 nearly-full-length (≥1560 nt) VP1 sequences from GII.4 strains collected from 1974 to 2016 (Table S1). The phylogenetic tree of these sequences showed the presence of at least 11 different GII.4 variants emerging since 1995 (Fig. 1a). As shown previously (15, 26), GII.4 strains presented a chronological replacement of variants, with several unassigned intermediate strains (Fig. S1). Genetic analyses revealed an accumulation of substitutions in both nucleotide (Fig. 1b) and amino acid (Fig. 1c) sequences (coefficient of determination [R^2^] of linear regression = 0.87 and 0.78, respectively). This pattern of accumulation of mutations was observed in all the subdomains of VP1 (Shell, P1, and P2; Fig. S2); however, higher slopes were noted in P2, where the five variable antigenic sites (A-E) are located, thus suggesting their role in the evolution and antigenic diversification of GII.4 noroviruses.

**Figure 1:**
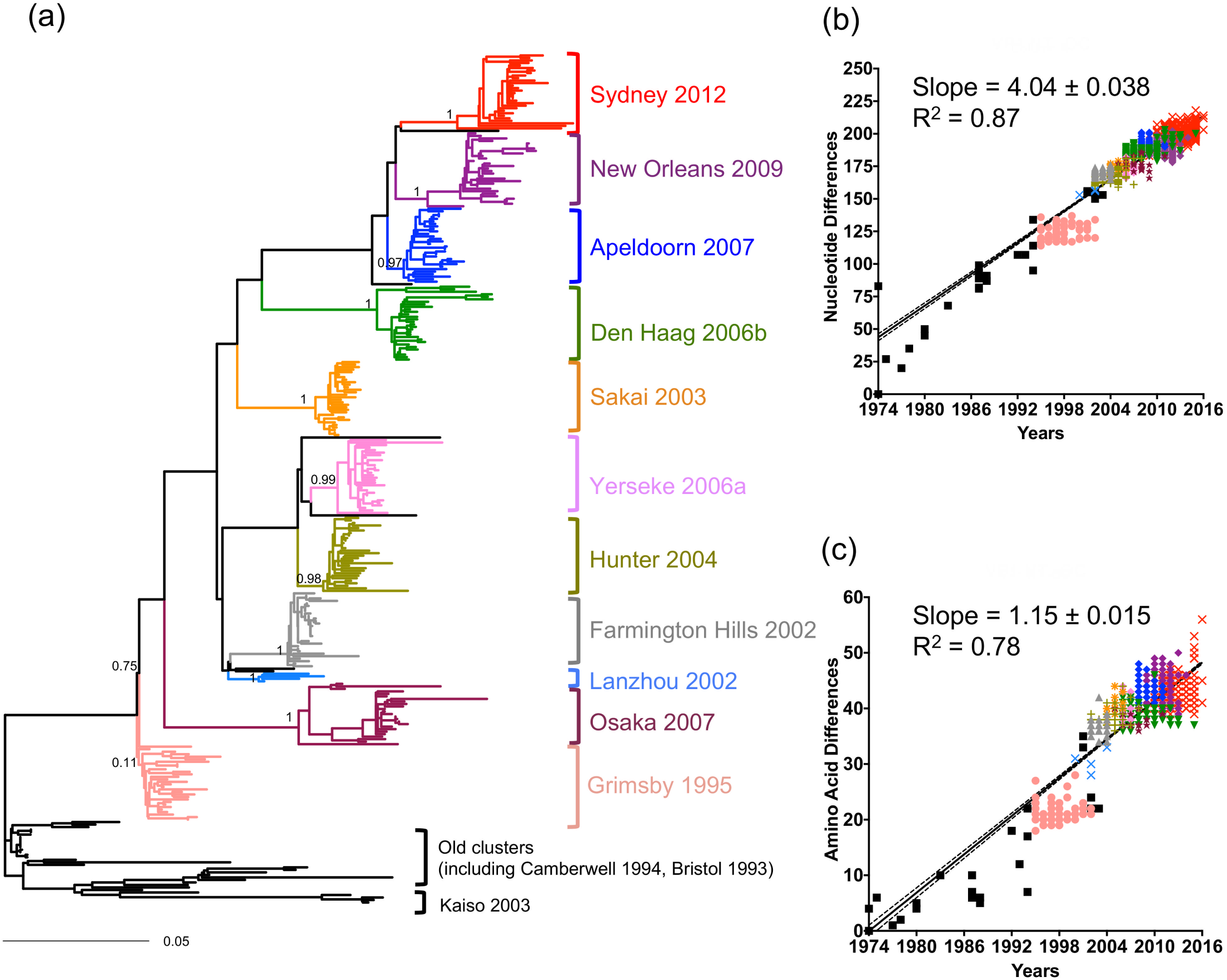
Evolution of the major capsid protein (VP1) from GII.4 noroviruses results in the accumulation of substitutions and the periodic emergence of variants. (a) Maximum likelihood tree of GII.4 noroviruses showing the circulation of different variants overtime. Branches are colored based on variant determination from the online Norovirus Typing Tool (https://www.rivm.nl/mpf/typingtool/norovirus/). Black branches in the tree represent sequences that did not cluster into a variant that circulated between 1995 and 2016. A subset of 308 sequences, from a total of 1601, were used for tree reconstruction as indicated in Materials and Methods section. Node support values calculated by the approximate likelihood-ratio test were shown on the major branches. Graphical representation of nucleotide (b) and amino acid (c) pairwise differences of each sequence in the dataset as compared to the earliest strain, a GII.4 strain collected in 1974 (AB303922). A total of 1601 nearly full-length ORF2 GII.4 sequences were included for the pairwise analyses. Black solid line indicates the linear regression line, with dotted lines showing the 95% confidence interval of the best-fit line.

### Identification of New Antigenic Sites of GII.4 Noroviruses

To pinpoint the role of each amino acid within the P domain in the evolution of GII.4, we calculated the Shannon Entropy to measure the residue diversity at the inter-variant and intra-variant level. Because Grimsby-like viruses were the first recorded to cause large outbreaks worldwide, entropy values were calculated with strains (1572 VP1 sequences) detected from 1995 to 2016. While minimal diversity was detected at the intra-variant level (Fig. S3), the inter-variant level revealed three substitution patterns: (i) a large number (87%, 285/325) of highly conserved residues, which include conserved antigenic site F (27), and likely maintain the structural integrity of the P domain; (ii) a small number (5%, 17/325) of variable residues that map on the previously determined antigenic sites: A-E (19); and (iii) 23 variable residues that map outside of those antigenic sites (Fig. 2a). Among the 23 non-antigenic variable sites from the third pattern, two residues (residues 228 and 255) were surrounded by conserved residues on the surface of VP1, and fourteen residues formed clusters (motifs) on the surface that could represent new, or extensions of previously predicted, uncharacterized antigenic sites (Fig. 2b). One motif comprises residues 339, 340, 341, 375, 376, 377, and 378. Because this motif included two residues (340 and 376) previously described as one of the five major antigenic sites (20), we extended the number of residues forming this antigenic site C (Fig. 2b). Although residue 375 presented low variability, the site was included as part of the antigenic site as it mapped on the surface of the molecule and could potentially play a role in antibody recognition. Antigenic site D was also extended to include two additional residues (396 and 397), as they clustered with original residues (393, 394, and 395) on the surface (19). The new motif, denominated G, included residues 352, 355, 356, 357, 359, and 364; and the last motif, denominated H, included the residues 309 and 310. A summary of the residues from each of the motifs/antigenic sites is shown in Fig. 2b. Profiling of the temporal frequency of the amino acid sequence pattern (mutational pattern) of the previously characterized antigenic site A, the expanded antigenic site C, and the new antigenic site G (confirmed in this study) indicated that their mutational patterns correlated well with the fluctuation of GII.4 variants in the human population, suggesting a major role in the emergence of variants (Fig. 3 and Fig. S4a). Correlation between the antigenic site mutational patterns and GII.4 variants was assessed using adjusted Rand Index, and antigenic site G was the one presenting the best correlation with the GII.4 variant circulation pattern, while showing low sequence variation (Fig. 3b and c). Of note is that the mutational pattern from the old antigenic site C did not correlate with the circulation of variants in nature (Fig. S4a and b), suggesting that our population-guided antigenic site characterization provides a better resolution of the antigenic profiling of GII.4 noroviruses. Neither the previously defined putative epitope (motif) B (19) nor the new variable motif H correlated with GII.4 variant circulation (Fig. 3c and Fig. S4a). Of note is that motif B presented differences for Farmington Hills 2002 and Hunter 2004 strains, while the motif H showed differences for Apeldoorn 2007, New Orleans 2009, and Sydney 2012 (Fig. S4a). Mutations on antigenic sites D and E have been shown to alter both antigenicity and binding to HBGA carbohydrates (19, 24, 28), while showing modest correlation with GII.4 variant emergence (Fig. 3c). Thus, antigenic sites D and E might be under different evolutionary pressures than those only involved in the antigenic characteristics of the virus. Of note, the mutational pattern from expanded antigenic site D correlates much better than that from the original antigenic site D (Fig. S4a and b), and this improvement of correlation is consistent with the recent addition of residue 396 to this antigenic site (29). On the contrary, the mutational pattern from the expanded antigenic site E showed lower correlation than that from the original antigenic site (Fig. S4a and b). This could be due to the expanded site (inclusion of residue 414) showing variation within the variants, Den Haag 2006b and Sydney 2012, making the profiles less consistent with the inter-variant diversity. Since our dataset presents sampling bias, i.e. over 50% of the sequences in the dataset were collected during 2010-2016, the Shannon entropy analysis was re-conducted using a randomly-subsampled dataset that includes a maximum of 50 sequences per variant (Fig. S5a). This dataset sensitively reflected the variation of viruses circulating before 2010, and showed 13 additional variable sites mapping outside the newly defined variable motifs/antigenic sites. Only three of them were located on the surface of the VP1; one of them (residue 250) did not cluster with any variable residues, and other two (residues 300 and 329) mapped close to the motif/antigenic site G (Fig. S5b). These two residues could be a part of the antigenic site G, however, both presented minor variation over decades suggesting a subtle impact on variant emergence (Fig. S5c).

**Figure 2:**
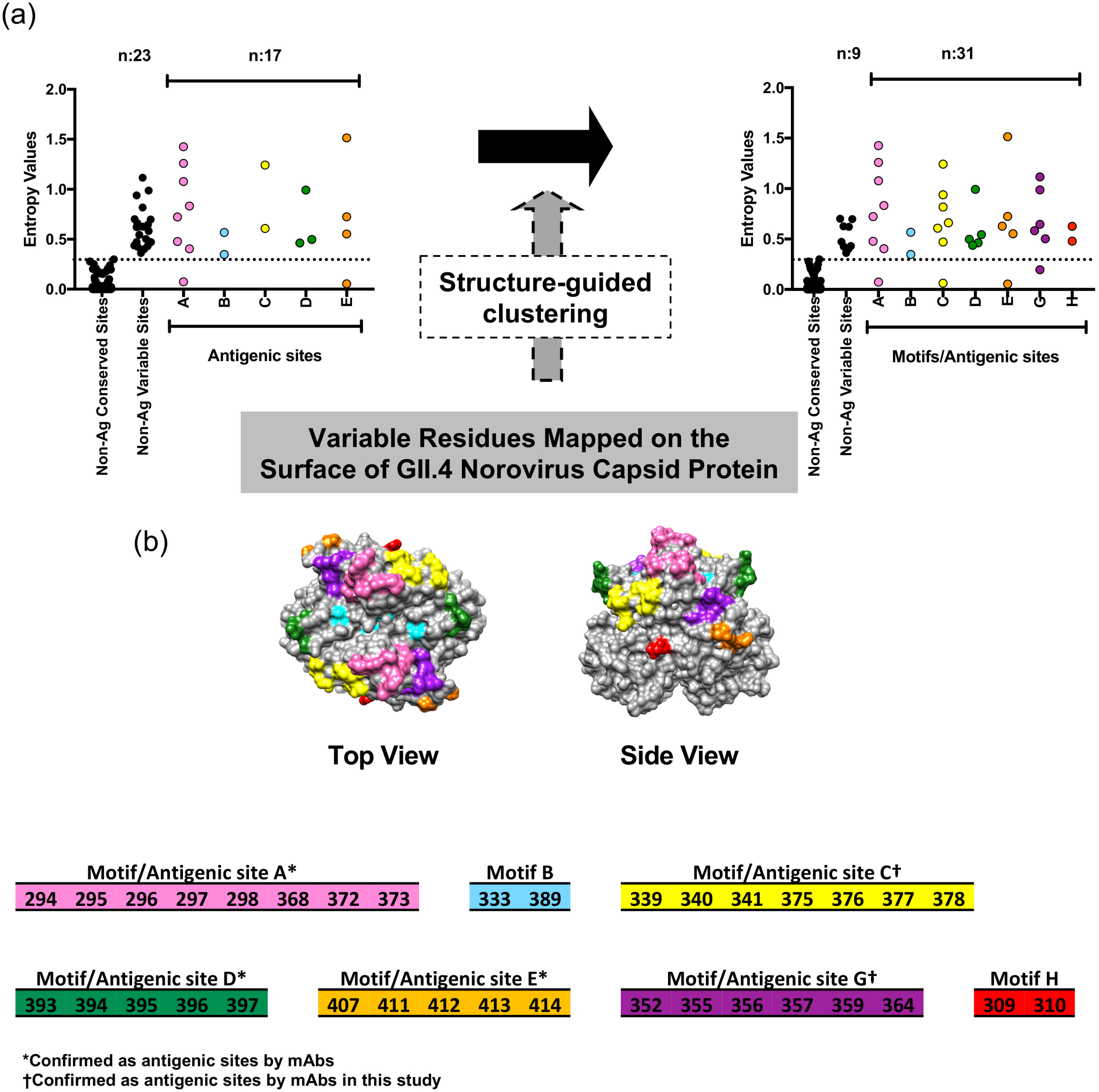
Conservation analyses redefined antigenic sites of the major capsid protein (VP1) from GII.4 noroviruses. (a) Shannon entropy was calculated to quantify amino acid variation for each site in the VP1. Analyses were calculated with strains (1572 VP1 sequences) detected from 1995 to 2016. Data from the P domain is included here (amino acids 216-540). Residues were grouped depending on the degree of variability into: conserved sites (Shannon entropy value ≤ 0.3), sites mapping on antigenic sites (A-E), and variable sites that map outside of antigenic sites (left-side dot plot). Based on structural analyses, 14 variable residues that mapped outside antigenic sites were clustered as part of novel motifs (potential antigenic sites) or extension of previously defined antigenic sites (right-side dot plot). (b) Residues forming these novel or expanded motifs/antigenic sites are colored accordingly on the surface of the GII.4 major capsid protein.

**Figure 3:**
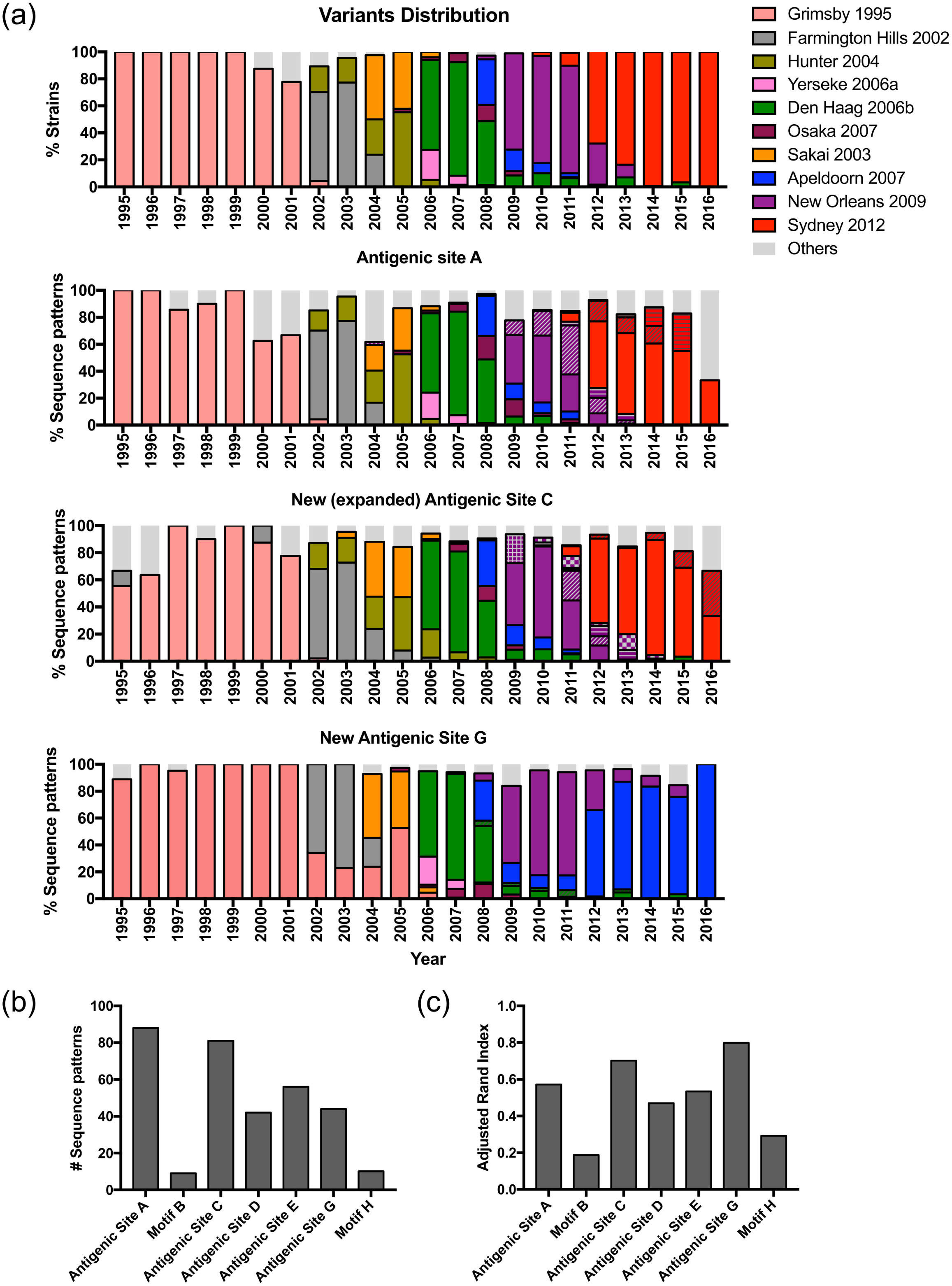
Predominant sequence patterns for proposed antigenic sites correlate with GII.4 variant circulation in the human population. **(a)** Amino acids from new and expanded motifs/antigenic sites were tabulated using 1572 sequences of GII.4 norovirus that circulated from 1995 to 2016. GII.4 variant yearly distribution was tabulated using the same sequence database. Colors of the bars for the profiling graphs correspond to the predominant sequence pattern presented in that antigenic site for each GII.4 variant. Colors and variant assignment follows those from Fig. 1. Patterns of the bars represent minor variations of the sequences in the motifs/antigenic sites. Motifs/Antigenic sites A, C, and G appear to follow the pattern of GII.4 variant distribution over time, implying the potential role of these sites in the emergence of new pandemic GII.4 variants. (b) Number of sequence patterns of each antigenic site and (c) the correlation between variant classification and mutational patterns of each motif/antigenic site were calculated. The degree of correlation was assessed by adjusted Rand Index, in which higher index indicates higher correlation between variant distribution and mutational pattern of the motif/antigenic site.

The mutational patterns from three motifs (A, C, and G) correlated well with the fluctuation of GII.4 variants (Fig. 3c). The motif/antigenic site A was previously confirmed as an antigenic site (23), while expanded/new motifs (C and G) were not yet confirmed experimentally. To confirm the role of these two motifs in the antigenic make-up of the GII.4 capsid, we replaced residues of the VP1 from a Farmington Hills 2002 variant (MD2004 strain; accession number: DQ658413) with those of a Sydney 2012 variant (RockvilleD1 strain; accession number: KY424328; Fig. 4) and vice versa, and produced the corresponding VLPs. We tested the mutant and wild-type VLPs against guinea pig hyper-immune sera and mouse mAbs (two uncharacterized mAbs: B11 and B12 raised against Farmington Hills 2002 variant (MD2004 strain) (11), and four mAbs: 1C10, 6E6, 17A5 and 18G12 newly developed against Sydney 2012 variant (RockvilleD1 strain)). We found that mutations at the new motifs C and G abrogated binding of mAbs B11, B12, 6E6, 18G12, and mAbs 1C10 and 17A5, respectively (Fig. 4a). Reconstitution of the motif C in the Sydney 2012 wild-type strain can be achieved by reverting those sites to Farmington Hills 2002 wild-type strain (2012: C2004 VLPs) (Fig. 4a, left panel). Of note, when mutations were introduced at residues 340 and 376, which were regarded as the original antigenic site C, no differences in binding were observed with mAb B11 and B12 (11). While changes at residues 377 and 378 reduced binding to those mAbs, an additional mutation at residue 340 was required for complete antigenic site-depletion (Fig. S6). Likewise, reconstitution of the motifs C and G in the Farmington Hills 2002 wild-type strain was achieved by reverting those sites to Sydney 2012 wild-type strain (2004: C2012 and 2004: G2012 VLPs) (Fig. 4a, right panel).

**Figure 4:**
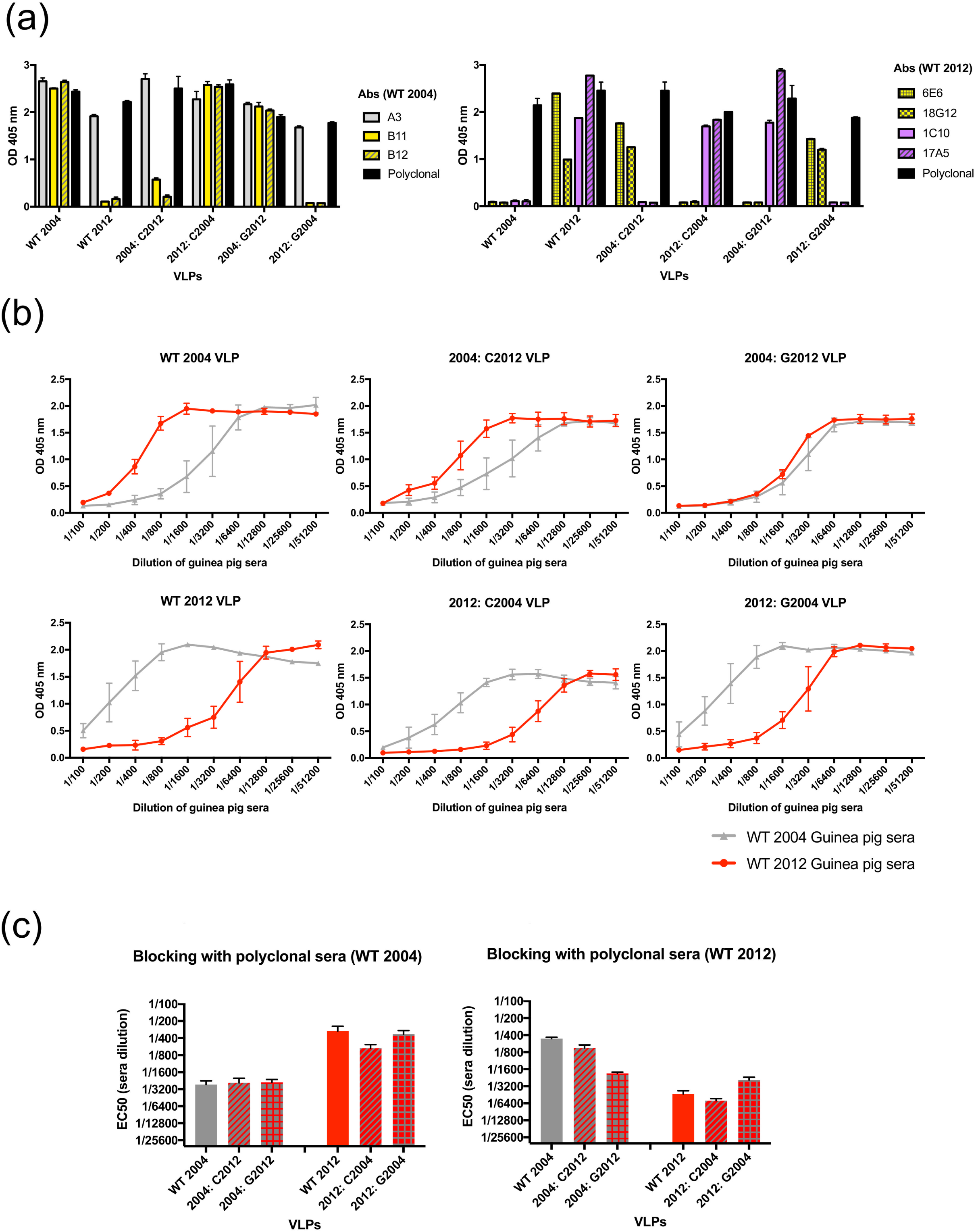
Mutational analyses on the major capsid protein (VP1) of GII.4 noroviruses confirmed newly proposed antigenic sites C and G. (a) Immunoassay performed with virus-like particles (VLPs) and mAbs raised against the Farmington Hills 2002 strain (MD2004-3 [DQ658413]; WT 2004 [(11)]) (left) and the Sydney 2012 strain (RockvilleD1 [KY424328]; WT 2012) (right). Mean and standard deviation (SD) were calculated from duplicate wells. (b) HBGA-blocking assays were performed with hyperimmune (polyclonal) sera raised against a Farmington Hills strain (MD2004-3; WT 2004) and a Sydney strain (RockvilleD1; WT 2012) and wild-type and mutant VLPs. Experiments were performed using sera from two guinea pigs for each strain, and mean and SD were calculated in duplicate wells. (c) The half-maximal binding (EC_50_) values (sera dilution) of each wild-type and mutant VLPs was calculated from the OD values from the blocking assays.

The impact of mutations on the motifs/antigenic sites C and G against the immune response was evaluated as blockade potential (a surrogate of norovirus neutralization) using HBGA-blocking assays. Polyclonal sera against Farmington Hills 2002 wild-type VLP showed high blocking activity against homologous wild-type VLP (low EC_50_ value), but reduced blocking against the Sydney 2012 wild-type VLP (high EC_50_ value) (Fig. 4b and c). Mutations on antigenic sites C and G on those wild-type VLPs altered the blockade potential of polyclonal sera. Thus, when antigenic site C from Farmington Hills 2002 wild-type strain was transplanted into Sydney 2012 wild-type VLP, polyclonal sera raised against Farmington Hills 2002 wild-type strain gained blocking potential against the Sydney 2012 mutant (2012: C2004) VLP, but minimally with the mutant 2012: G2004. Notably, none of the Farmington Hills 2002 mutant VLPs (2004: C2012 and 2004: G2012) lost the blocking ability when using sera raised against the Farmington Hills strain. Similarly, polyclonal sera raised against Sydney 2012 wild-type VLPs blocked the homologous VLPs, but not the Farmington Hills 2002 wild-type VLPs. Transplanting of antigenic site C and G into Farmington Hills 2002 VLPs (2004: C2012 and 2004: G2012) recovered its blocking ability of sera raised against Sydney 2012 wild-type strain. Polyclonal sera raised against Sydney 2012 wild-type VLPs did not lose blocking activity against antigenic site C mutant VLP (2012: C2004), however, a slight lost in the blocking activity was detected for antigenic site G mutant (2012: G2004) as compared with wild-type VLP. In summary, both antigenic sites (C and G) showed distinctive roles as blockade sites, suggesting their potential as protective antigenic sites.

### Intra-variant Evolution is Driven by Stochastic Processes

In contrast to the accumulation of nucleotides and amino acids detected at the inter-variant level (Fig. 1b and 1c), there were limited accumulation of nucleotide (data not shown) and amino acid substitutions within variants (intra-variants level) (Fig. 5a). Despite this limited accumulation of substitutions, each of the pandemic variant presented diversity in their sequences (average of 4.8 amino acid substitutions). Interestingly, this diversity was detected at most of the major antigenic sites and variants (Fig. 5b and Fig. S7). Thus, while each variant presented a major amino acid combination for each antigenic site, most of the GII.4 variants presented other minor amino acid patterns on those antigenic sites. Moreover, the analysis of intra-variant diversification showed that their evolution was stochastic in time and location (Fig. 6), in contrast to the temporally clustered inter-variant evolution of GII.4 strains (Fig. 1). While some variants (e.g. Hunter 2004, Den Haag 2006b, New Orleans 2009, and Sydney 2012) presented diversity in their major antigenic sites after 3-4 years of circulation, which could suggest that immune pressure acts at the intra-variant levels (Fig. 5), the number of strains (and sequences) is limited and do not represent dominant strains. Two other important observations could be made while analyzing the mutational pattern of each of the antigenic sites at the intra-variant level: (i) major differences at the amino acid sequence were detected in early strains for New Orleans 2009 and Sydney 2012 variants (Fig. 5 and Fig. S7), and those early strains did not present the major amino acid combination for any of the four major antigenic sites (A, C, D, and E; Fig. S7); and (ii) none of the preceding strains evolved towards (or presented) the amino acid motif from future strains. This, together with the phylogenetic analyses, suggests that each pandemic variant presented a different origin and did not follow a trunk-like linear evolution such as that seen in H3N2 influenza viruses (30).

**Figure 5:**
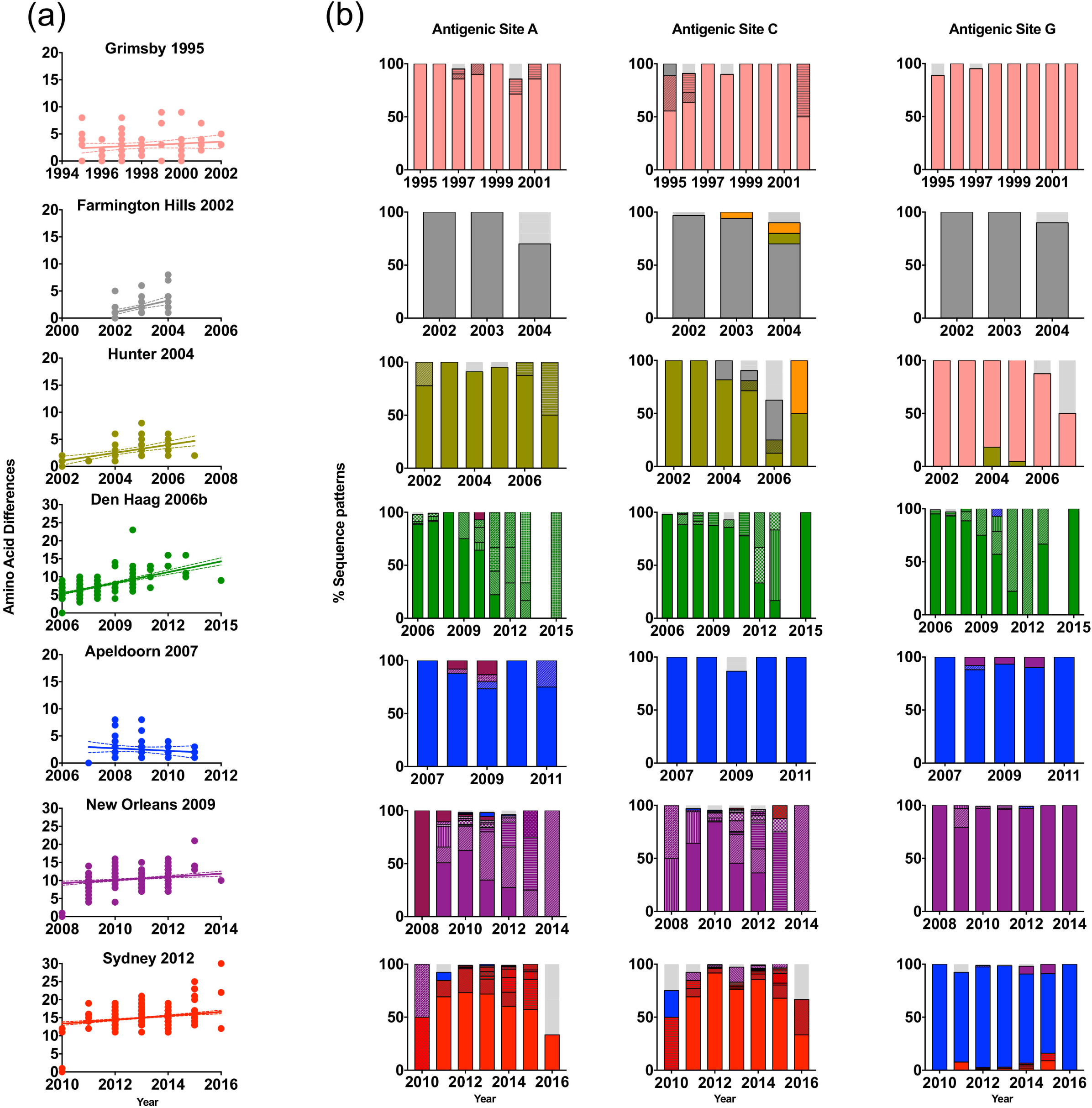
Intra-variant diversity of GII.4 noroviruses reveals minimal accumulation of mutations. (a) Amino acid pairwise differences among viruses from each of the pandemic GII.4 variants as compared to the earliest strain from each given variant. (b) Intra-variant mutational pattern of each antigenic site for each pandemic GII.4 variant was calculated as indicated in Fig. 3.

**Figure 6:**
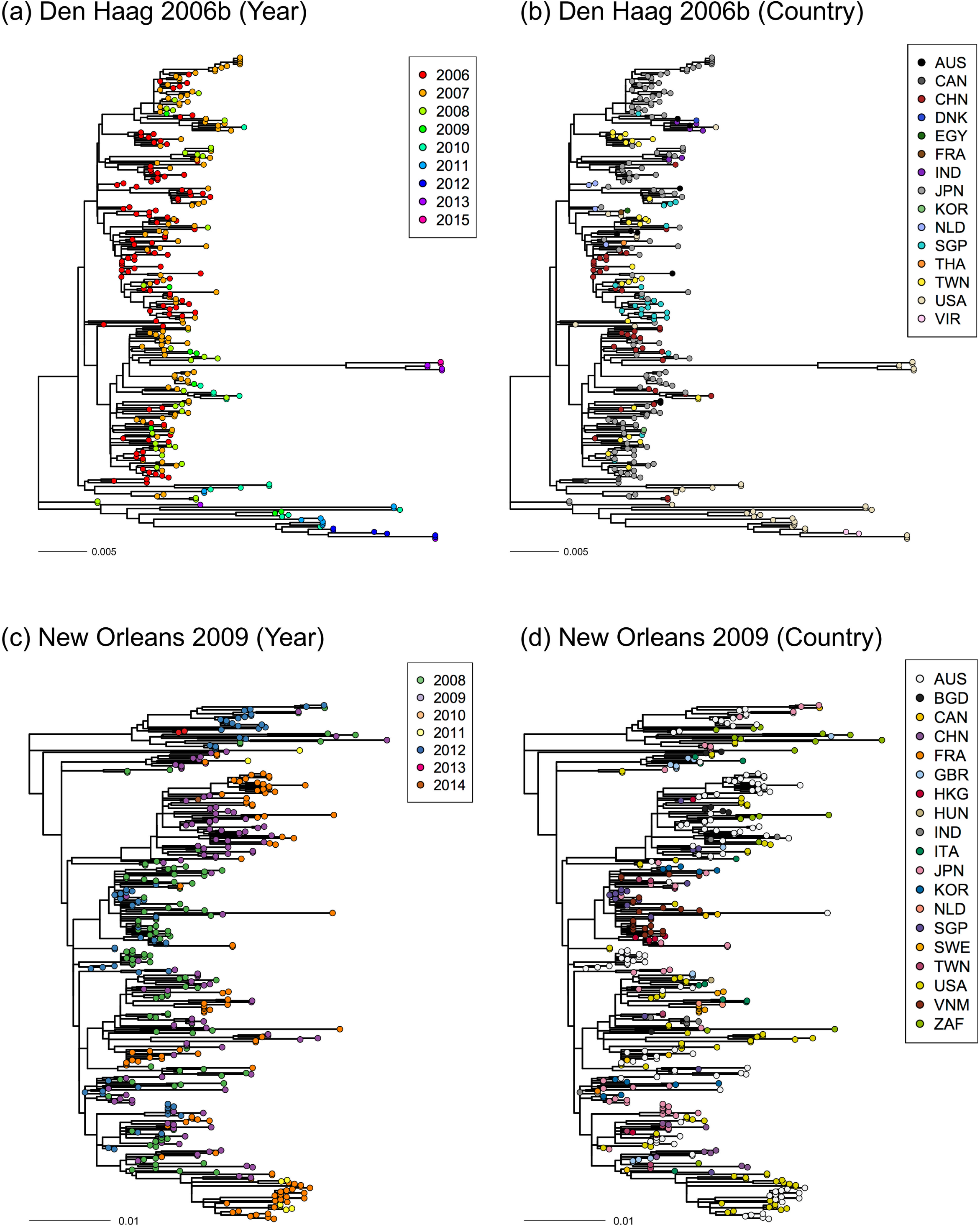
Intra-variant diversification of GII.4 noroviruses is governed by stochastic events. Maximum likelihood trees of two major GII.4 variants, Den Haag 2006b and New Orleans 2009, showing the collection year (a and c) or country (b and d) for each of their strains. Phylogenetic clustering of the strains did not present any pattern indicating randomness of the intra-variant evolution in time and space.

Differences in the inter- and intra-variant evolutionary pattern were also confirmed by the Bayesian Markov Chain Monte Carlo (MCMC) analysis. Substitution rates from seven variants with > 50 sequences were calculated and summarized in Table 1. The rate of inter-variant (overall) GII.4 strains was reported elsewhere (15). Intra-variant substitution rates ranged from 1.57 – 4.64 × 10^-3^ substitutions/site/year, all of them lower than the inter-variant (overall) substitution rate (5.4 × 10^-3^ substitutions/site/year) (15). Among the variants, Sydney 2012 variants had the higher substitution rate (Table 1).

**Table 1:**
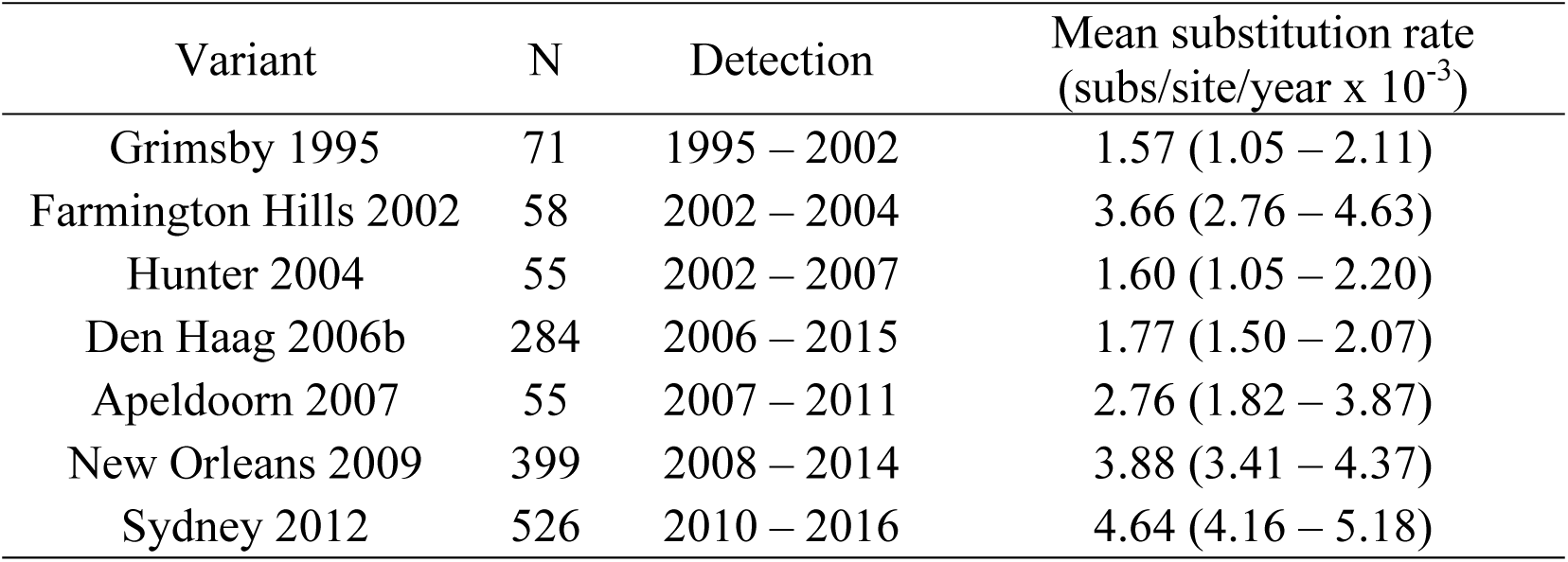
GII.4 variants rate of evolution

| Variant | N | Detection | Mean substitution rate<br>(subs/site/year x 10<sup>-3</sup>) |
| --- | --- | --- | --- |
| Grimsby 1995 | 71 | 1995 – 2002 | 1.57 (1.05 – 2.11) |
| Farmington Hills 2002 | 58 | 2002 – 2004 | 3.66 (2.76 – 4.63) |
| Hunter 2004 | 55 | 2002 – 2007 | 1.60 (1.05 – 2.20) |
| Den Haag 2006b | 284 | 2006 – 2015 | 1.77 (1.50 – 2.07) |
| Apeldoorn 2007 | 55 | 2007 – 2011 | 2.76 (1.82 – 3.87) |
| New Orleans 2009 | 399 | 2008 – 2014 | 3.88 (3.41 – 4.37) |
| Sydney 2012 | 526 | 2010 – 2016 | 4.64 (4.16 – 5.18) |

### Diversifying Pressure Drives Emergence of Pandemic GII.4 Noroviruses

Because different factors seemed to drive the inter-variant (overall) and intra-variant evolution, we performed selection analysis using the Mixed Effect Model of Evolution (MEME) method and looked for evidence of site-by-site episodic diversifying pressure on the VP1 along the branches of its evolutionary tree. To analyze the overall evolutionary process, we randomly subsampled sequences from the original dataset that included a maximum of 30 strains per variant. During the overall evolution, we found 9 positively selected sites (*p* < 0.05 and empirical Bayes factor > 100) on the P2 subdomain on the surface (codon sites: 327, 335, 352, 355, 357, 366, 368, 375, and 378) distributed on 21 branches of the phylogenetic tree (Fig. 7a). Branches connecting discrete GII.4 variants presented sites under episodic diversification (codon sites: 352, 355, 357, 368, 378) that mapped on the antigenic sites. The mutational pattern of these sites correlated well with GII.4 variant emergence, with higher adjusted Rand Index than any antigenic sites (Fig. 7b and c). This suggests that residues on the antigenic sites experienced episodic diversifying pressure during inter-variant evolution. When analyzing only each variant, most of the diversifying pressure was found on the branches connecting to the tips rather than on the internal branches (Fig. S8), suggesting that non-synonymous substitutions were deleterious and rarely fixed in the population during the intra-variant evolution. The more comprehensive dataset presented by New Orleans and Sydney variants may have included many viruses with deleterious mutations that would not persist (as indicated by higher number of non-synonymous substitutions on branches connecting to the tips), but would account for an artificial higher substitution rate as compared with the other variants (Table 1). Together, this shows that the diversifying pressure has driven the inter-variant, but not the intra-variant, evolution of GII.4 strains.

**Figure 7:**
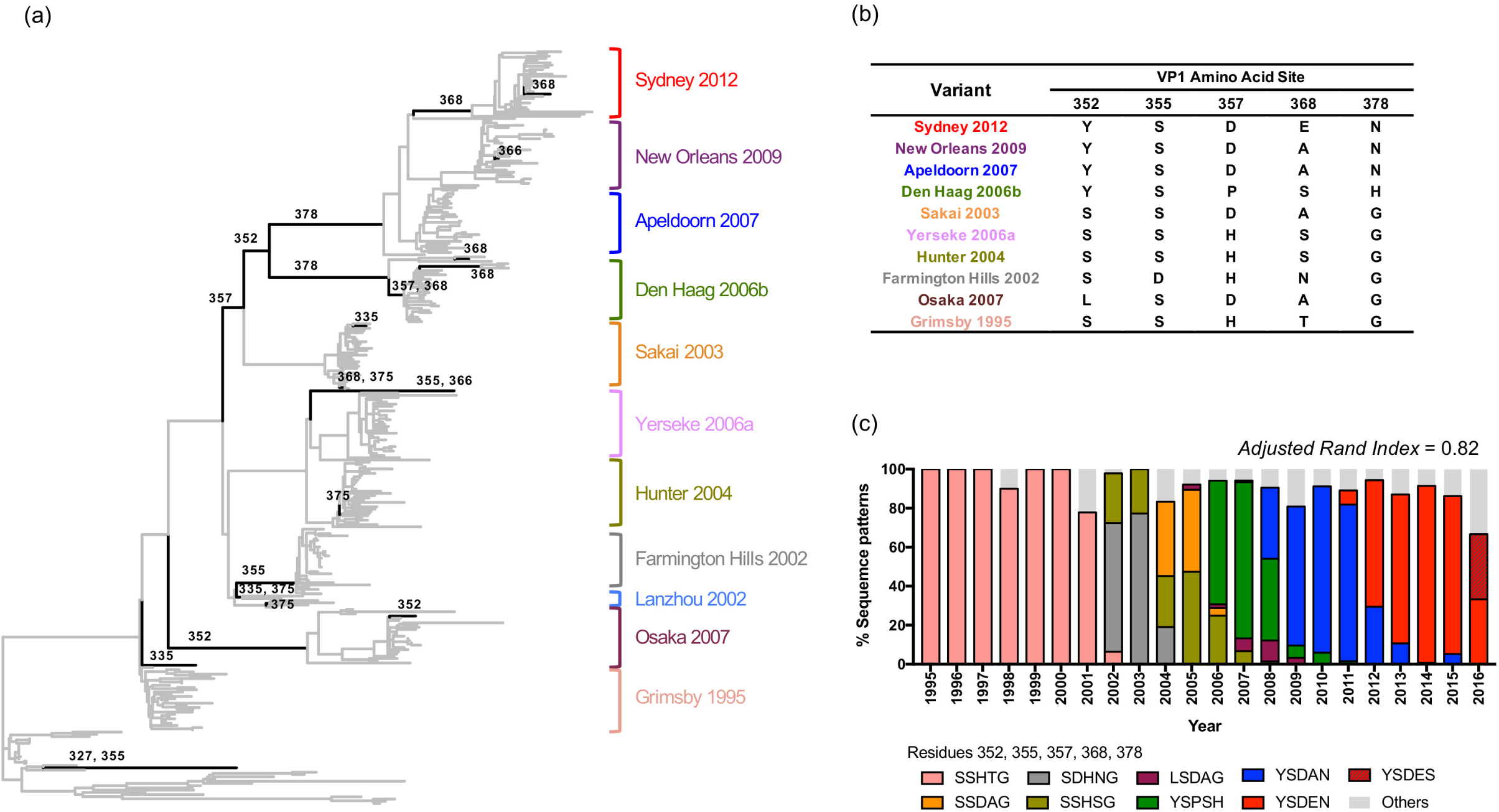
Diversifying pressure on GII.4 evolution. (a) Maximum likelihood tree of all GII.4 variants indicates the branches under possible diversifying selection. The branches under diversifying selection (empirical Bayes factor > 100 and *p* < 0.05) were explored using Mixed Effect Model of Evolution method and codon positions that are located on the capsid surface are represented by black branches. Diversifying selection at codon positions 352, 355, 357, 368, and 378 appeared on the inter-variant branches, suggesting their significant role in the emergence of new GII.4 variants. (b) Predominant amino acids present in each of the sites under diversifying selection for each GII.4 variant that emerged since 1995. (c) The mutational patterns and adjusted Rand Index for the sites under diversifying selection were calculated as indicated in Fig. 3.

## Discussion

GII.4 noroviruses are the most common cause of norovirus infections worldwide. Although other norovirus genotypes have predominated in specific locations and time, the global dominance of GII.4 has been recorded for almost three decades (16, 31). The persistence and dominance of GII.4 over all other norovirus genotypes has been explained by the chronological emergence of variants, which enables the virus to evade the immunity acquired to previously-circulating GII.4 variants, a process similar to H3N2 influenza viruses (26, 28, 30). Since the mid-1990s, over 10 different variants have been reported, with six of them associated with large outbreaks worldwide. The overall evolutionary pattern of GII.4 viruses presents a strong linear accumulation of amino acid substitutions during inter-variant diversification (15), with most substitutions occurring in the P2 subdomain (28). Antigenic differences among variants have been largely attributed to highly variable residues that map on the surface of the P domain, leading to the identification of five (A-E) motifs that are part of GII.4-specific antigenic sites (18–20). While the binding site was characterized for different GII.4 specific mAbs, the same studies have reported numerous GII.4 specific mAbs whose binding sites have not been determined (11, 12, 19, 23–25). Applying a population genomics approach we found new (or expanded) motifs on the surface of the capsid that presented mutational patterns that correlates with the circulation of GII.4 variants. The role of antigenicity of two of those motifs (antigenic sites C and G) were confirmed with (previously) uncharacterized HBGA-blockade mAbs (B11 and B12), newly developed mAbs (1C10, 6E6, 17A5, and 18G12), and polyclonal sera from guinea pigs immunized with wild-type VLPs. The antigenic site G presented the strongest correlation with the emergence and circulation of new GII.4 variants, while being more conserved when compared with other large antigenic sites (e.g. A, C, or E). This indicates that newly discovered antigenic site G plays a pivotal role in shaping the GII.4 noroviruses into new variants with pandemic potential. The antigenic site C is close to the previously-defined antigenic site A, but substitutions on antigenic site A did not affect the binding of those mAbs (11). Notably, competition analyses among different mAbs showed that mAbs B11 and B12 partially blocked interactions with mAbs mapping to antigenic site A (11), suggesting that this new motif is part of an antigenic site that involves more than one epitope (at least antigenic sites A and C). Recently, Koromyslova and colleagues have shown that the footprint of an antibody that neutralized human noroviruses mapped to residues from antigenic sites C and D (22). This demonstrates that same epitope could be shared by the different antigenic sites and their interaction could result in differences in the evolutionary pattern presented. In addition, differences on the HBGA-blocking ability between antigenic sites (C and G) and between variants (Farmington Hills 2002 and Sydney 2012) further highlight the complexity of the antigenic topology of GII.4 noroviruses.

Large-scale analysis reduces biases on the role of individual amino acid substitution on the emergence of the new GII.4 variants. Recently, it was suggested that mutations in the capsid protein of Sydney strains circulating in 2015 resulted in antigenically different viruses from those circulating in 2012 (32). Our large-scale intra-variant analyses show that (i) the strain selected as Sydney 2012 for the antigenic study (32) was not representative of the predominant virus, and (ii) strains with similar antigenic site sequences as those regarded as “GII.4 2015” by Lindesmith et al. (32) have been circulating since 2010 in the human population as part of the overall population of the Sydney 2012 variant (Fig. S7). Similarly, the NERK motif, which was suggested to occlude conserved GII.4 antigenic sites and effect the antibody blockade potency (33, 34), was shown to be highly conserved among GII.4 variants. Previous studies pointed out a mutation on this motif in Sydney 2012 variant, however, our large-scale analysis showed that the New Orleans variant was the only one showing -at the population level-any mutation at this motif (Fig. S9). Thus, the profiling of mutational patterns we implemented, which included the use of a large number of sequences, could provide a better understanding on the role of individual mutations on the circulation and predominance of the pandemic GII.4 variants. Immunological analyses that include multiple different viruses from each of the pandemic variants are needed to better delineate the meaning of minor mutations on the antigenic differences among the variants.

Improvements on the understanding of viral dynamics and the correlation between antigenic and genetic changes in influenza viruses have facilitated the selection of virus strains to be included in upcoming seasonal vaccines (35–37). To better understand viral dynamics of GII.4 noroviruses, we performed selection analyses including all variants reported for over four decades. Episodic diversifying (positive) selection was observed in five residues (352, 355, 357, 368, and 378) during inter-variant evolution (Fig. 7a); these residues are part of antigenic sites A, C, and G. Three of these residues (352, 357, and 378) showed positive selection on major branches of the GII.4 tree, indicating a role in the emergence of new pandemic noroviruses. In these three sites, strains that emerged and predominated from 1995 to 2006 (Grimsby 1995, Farmington Hills 2002, Hunter 2004, and Yerseke 2006a) presented the motif SHG; the strain Den Haag 2006b that emerged and predominated from 2006 to 2009 presented the motif YPH; the strains that have predominated since 2009 (Apeldoorn 2007, New Orleans 2009, and Sydney 2012) presented the motif YDN. Notably, while these three residues have been understudied pertaining to the evolution of GII.4 noroviruses (26, 28, 38), they seem to play a major role in the antigenic topology and emergence of new GII.4 variants. Of note, residue 368 presented episodic diversification that distinguished the Sydney 2012 strains from Apeldoorn 2007 and New Orleans 2009. Changes on this residue were shown to be involved in the antigenic diversification of the Sydney 2012 strain (39), thus supporting our *in silico* observation. Initial studies suggested that most positively-selected sites for GII.4 noroviruses were located in the S domain (28, 38, 40); however, our analyses, with an expanded number of sequences and variants, showed that the emergence of new variants stemmed from the positive selection of residues mapping on the surface of the P domain. Pinpointing the changes required for emergence of a new antigenic variant of seasonal influenza or the emergence of a pandemic virus is the “holy grail” for controlling viruses undergoing constant change. While prediction of viral emergence requires a wholistic approach that includes studies of the virus, environment, and the host (41–43), careful monitoring of the substitutions in residues involved in the diversification and emergence of new GII.4 viruses (i.e. residues 352, 357, and 378) could help in the early detection of future novel variants with pandemic potential.

The GII.4 inter-variant diversification, likely driven by the immune status of the population, is correlated with the accumulation of amino acid substitutions in the major capsid protein. In contrast, our analyses suggest that the diversification at intra-variant level is much restricted, with amino acid substitutions occurring without indication of diversifying selection. Minimal substitutions of the amino acids that mapped on the major antigenic sites were observed over the predominance of each variant, and most seemed to follow a stochastic process. The latter could be a result of multiple pressures on the virus, including but not limited to individual virus-host interactions and dispersion. For two variants (New Orleans 2009 and Sydney 2012) a larger, more comprehensive dataset was available, and pre-pandemic strains have been reported (44–47). Interestingly, although these pre-pandemic strains cluster within their respective variants, they present multiple differences (mostly mapping at the antigenic sites), as compared to the virus population that later established a worldwide dominance. Together, these results suggest that emergence of GII.4 variants could occur in steps: (1) a pre-pandemic stage characterized by acquisition of mutations that facilitate viral emergence and episodic diversification (exemplified here by residues 352, 357, 368, and 378), followed by (2) a short period (1-2 years) of adaptability (with different antigenic motifs) that precedes the pandemic phase, and finally (3) the pandemic phase, where the virus is dominant and only explores a narrow space of sequence diversity. A similar pattern has been observed for H3N2 influenza viruses (48) or rotaviruses (49), in which viruses that circulate at low levels could predominate in the upcoming season without major changes in their genetic background. This order of events could have also occurred during the recent emergence and predominance of GII.17 noroviruses in many Asian countries. During 2013-2014, a “new” GII.17 (variant C) was detected circulating in different countries (Japan, Hong Kong, China), but in the next epidemic season, this variant was ultimately replaced by the variant D that spread worldwide and predominated from 2014-2016 (15, 50–52). In the same context, we found six GII.4 strains that did not cluster with any GII.4 variant (Fig. S1), and could represent strains that did not adapt well to the human population. These strains presented different sequence patterns in their antigenic sites, and therefore could constitute strains in the pre-pandemic stage that explored the sequence space but failed to thrive to the next evolutionary level.

Our findings suggest two different mechanisms behind the evolutionary dynamics of the major capsid protein from GII.4 noroviruses: (1) a steady pandemic phase governed by stochastic processes, which is preceded by (2) substitutions that arise from positive selection. Our study also provides a methodological framework that could facilitate the characterization of variable antigenic sites that play a relevant role in the emergence of new viruses in the human population. Studies that include large and long time-scale datasets of full-length genomes would help to determine the factors involved in norovirus predominance and persistence in the human population.

## Materials and Methods

### 1. Data mining and sequence analyses

A total of 1601 full-length (1623 nt) and nearly full-length (≥1560 nt) VP1 sequences of the GII.4 genotype, in which sequences from immunocompromised patients and environmental samples were removed, were downloaded from GenBank (accessed July 2017) (Table S1). The sequences spanned 42 years from 1974 to 2016. Sequences were aligned using ClustalW as implemented in MEGA v7 (53), and visually inspected to confirm proper alignment. Nomenclature for GII.4 variants was adopted as previously indicated (15), and variant datasets were parsed following such information. Sequence analyses were performed using total dataset except where indicated. Entropy analysis and profiling of mutational patterns only included strains (1572 VP1 sequences) collected from 1995 to 2016 as Grimsby-like viruses were the first recorded to cause large outbreaks worldwide (31), and included the following 11 variants in approximate order of emergence: Grimsby 1995 (or US95_96), Farmington Hills 2002, Lanzhou 2002, Sakai 2003 (or Asia 2003), Hunter 2004, Yerseke 2006a, Den Haag 2006b, Osaka 2007, Apeldoorn 2007, New Orleans 2009, and Sydney 2012. Intra-variant and inter-variant Shannon entropy was calculated using the Shannon Entropy-One tool as implemented in Los Alamos National Laboratory (www.hiv.lanl.gov). Entropy values for each position were plotted in GraphPad Prism v7. The structural model of GII.4 norovirus P domain dimer (Protein Data Base [PDB] accession number: 2OBS) was rendered using UCSF Chimera (version 1.11.2) (54). Profiling of mutational patterns of motifs/antigenic sites was performed using R 3.4.2 (55). Amino acid sequences of each motif/antigenic site was profiled by year and its mutational patterns were plotted in a composite bar graph showing the number of strains with each pattern as a fraction (%) of a whole (total number of strains). The correlation of the mutational patterns and variant distribution was assessed using adjusted Rand Index, known as a clustering analysis method, which evaluated the degree of the matches between mutational patterns and variant classification. Adjusted Rand Index was calculated using R and *mclust* package (56). To account for the sampling bias from the original dataset, we repeated entropy and structural analysis using randomly subsampled dataset that included a maximum 50 strains/variant (n = 474) from the original GII.4 dataset (1572 sequences).

### 2. Diversifying selection analysis

Maximum likelihood (ML) phylogenetic trees of VP1-enconding nucleotide sequences for all variants (inter-variant analyses) and each variant (intra-variant analyses) were constructed using the PhyML (57). The best substitution models were selected based on the lowest Akaike Information Criterion (AICc) for each dataset using jModelTest v2 (58, 59). Larger datasets (i.e. Sydney 2012 and New Orleans 2009 variants) tended to favor a Generalized time-reversible (GTR) substitution model, while variants with fewer reported sequences favored a Tamura-Nei 93 (TN93) model. Diversifying selection on the VP1-encoding sequence through its ML phylogenetic tree was analyzed by using MEME methods (60, 61). We aimed to detect codon sites under positive selection (i.e. more non-synonymous substitutions than synonymous substitutions) during the evolution and focused on the sites on or near the major antigenic sites of GII.4. Significant positive selection was indicated by *p* < 0.05. The branches under diversifying positive selection were explored by using empirical Bayes factor >100 in MEME. To reduce the data size for the inter-variant analyses of positive selection, we randomly subsampled a maximum 30 strains/variant (n = 308) from the original GII.4 dataset (1601 sequences) as indicated previously.

### 3. Bayesian analyses of nucleotide substitution rate

Using the VP1 sequences and respective collection years from each strain, temporal phylogenetic analysis was calculated using Bayesian Markov Chain Monte Carlo (MCMC) methodology in BEAST v1.8.3 (62). The best substitution models were selected based on the lowest Akaike Information Criterion (AICc) as mentioned above. The clock models (strict or relaxed lognormal clock) and tree priors (constant population size, exponential growth, or skyline) were tested, and best models were selected based on the model selection procedure using AIC through MCMC. The MCMC runs were performed until all the parameters reached convergence. MCMC runs were analyzed using Tracer v1.6 (http://tree.bio.ed.ac.uk/software/tracer/). The initial 10% of the logs from the MCMC run were removed before summarizing the mean and the 95% highest posterior density interval of the substitution rates.

### 4. Site-directed Mutagenesis and VLPs production

The VP1-encoding sequences from a GII.4 Farmington Hills 2002 (MD2004-3) and Sydney 2012 (RockvilleD1) strain were ligated into pFastBac1 vectors using SalI and NotI restriction sites (11, 63). Site-directed mutagenesis of pFastBac-MD2004-3 and pFastBac-RockvilleD1 was performed using mutation-specific forward and reverse primers, followed by purification with illustra MicroSpin G-50 Columns (GE Healthcare, Buckinghamshire, United Kingdom). Parental DNA was digested using the Dpn1 enzyme (New England BioLabs, Massachusetts, USA). VLPs presenting multiple mutations were developed by cloning a chemically synthesized P domain into a pFastBac1 plasmid containing the S domain using PspXI and NotI restriction sites (11). Each pFastBac construct was transformed via electroporation into ElectroMAX DH10B Cells (Thermo Fisher Scientific, California, USA), and grown on LB plates with ampicillin overnight at 37°C. Selected colonies were used to extract plasmid DNA (QIAprep Spin Miniprep Kit; Qiagen, Hilden, Germany). Introduction of mutations were confirmed by Sanger sequencing. VLPs were produced using the Bac-to-Bac Baculovirus Expression System (Invitrogen, California, USA) and purified through a cesium chloride gradient as previously described (63). Expression of VP1 protein was confirmed by western blot, and VLP integrity was confirmed by electron microscopy.

### 5. Immunoassays

Mutants and wild-type norovirus VLPs were analyzed for reactivity to monoclonal antibodies by enzyme-linked immunosorbent assay (ELISA) as described previously (11). The mAbs B11 and B12 were obtained from mice immunized with GII.4 Farmington Hills 2002 variant (MD2004-3 strain) VLP (11), and generously provided by Dr. Kim Y. Green (National Institutes of Health, USA). The GII.4 Sydney 2012 variant specific-mAbs (1C10, 6E6, 17A5, and 18G12) were developed from mice immunized with RockvilleD1 strain VLP (GenScript, NJ, USA). HBGA-blocking assays were performed using mutant and wild-type VLPs, HBGA molecules derived from human saliva, and polyclonal antibodies. The polyclonal antibodies were obtained from guinea pigs and mice immunized with GII.4 Farmington Hills 2002 (MD2004-3) and Sydney 2012 (RockvilleD1 strain) VLPs (11, 63). Human saliva was collected from a healthy adult volunteer. Saliva sample was boiled at 100°C for 10 mins immediately after the collection, and centrifuged at 13,000 rpm for 5min. The saliva supernatant was collected and used for HBGA-binding and -blocking assays. Briefly, serial dilution of guinea pig polyclonal sera were mixed with wild-type or mutant VLPs and incubated on the saliva-coated plate for 1h at 37°C. Plates were washed four times to remove unbound (i.e. blocked by guinea pig polyclonal sera) VLPs. Pooled sera from mice immunized with Farmington Hills 2002 VLP and Sydney 2012 VLP was used to detect the VLPs attached to the plate. Goat anti-mouse IgG conjugated with horseradish peroxidase and 2,2’-azino-bis(3-ethylbenzothiazoline-6-sulphonic acid) (ABTS) substrate (SeraCare, MA, USA) were used to develop the blue-green color on the VLP-attached plate. The degree of blocking was evaluated using optical density (OD) at 405 nm and EC_50_ of sera dilution. The EC_50_ was calculated from the OD curve using GraphPad Prism v7.

## Acknowledgments

We thank Dr. Steve Rubin for the critical reading of the manuscript. Financial support for this work was provided by the Food and Drug Administration intramural funds [Program Number Z01 BK 04012-01 LHV to G.I.P]. K.T and C.J.L are recipients of a CBER/FDA-sponsored Oak Ridge Institute for Science and Education (ORISE) fellowship.

## Supplemental Material

**Supplementary Figure 1: Unassigned strains which could not be clustered with any variants in the phylogenetic tree.** (a) Six strains which could not be assigned and located outside the variant clusters were indicated on the Maximum likelihood tree of GII.4 noroviruses. Branches are colored based on variant determination as in Fig. 1. The table shows amino acid sequences of the variable motifs/antigenic sites of those unassigned strains. The color in the table corresponds to the predominant pattern presented in that antigenic site for each GII.4 variant.

**Supplementary Figure 2: Pairwise differences of GII.4 VP1 sequence database indicated by the structural subdomains.** (a) The structural model of norovirus VP1 (PDB accession number: 1IHM) was rendered using UCSF Chimera (version 1.11.2). (b-d) Pairwise differences were calculated and plotted as in Fig. 1b, except that sequences spanning each of the individual structural subdomains of VP1 were used for the analyses. The structural subdomains of norovirus VP1 are defined as the following: P2 (b), amino acids 281-415; P1 (c), N-terminal amino acids 216-280, C-terminal amino acids 416-540; and Shell (d), amino acids 1-215.

**Supplementary Figure 3: Conservation analyses of the major capsid protein (VP1) from GII.4 noroviruses.** Shannon entropy values were calculated to quantify amino acid variation for each site in the VP1. The top panel presents Shannon entropy values for the P domain for all GII.4 sequences available in public databases. The bottom panels present Shannon entropy values for the P domain of each of the seven major GII.4 variants that emerged since 1995. Sites mapping at the variable motifs/antigenic sites (A-E, G, and H) are indicated by different colors.

**Supplementary Figure 4: Amino acid mutational patterns comparing the new and old variable motifs/antigenic sites.** (a) Mutational patterns of all the variable motifs/antigenic sites (A-E, G, and H) and the previously defined original antigenic sites C (amino acids 340, 376), D (amino acids 393-395), and E (amino acids 407, 411-413). GII.4 variant distribution was plotted as described in Fig. 3. Amino acids from each new and expanded motifs/antigenic sites were tabulated using 1572 sequences of GII.4 norovirus that circulated from 1995 to 2016. Colors of the bars for the profiling graphs correspond to the predominant sequence pattern presented in that motif/antigenic site for each GII.4 variant. Patterns of the bars represent minor variations of sequences in the motifs/antigenic sites. The amino acid sequence patterns of each motif/antigenic site were listed in the legend of each bar graph. (b) Adjusted Rand Index shows higher correlation of mutational pattern of expanded antigenic site C and D compared with that of the original antigenic site C and D, respectively. Mutational patterns of expanded antigenic site E was less correlated with variant distribution as compared with that of the original antigenic site E.

**Supplementary Figure 5: Conservation analyses of the major capsid protein (VP1) from a randomly subsampled dataset.** (a) To account for sampling bias, Shannon entropy was re-calculated to quantify amino acid variation for each site in the VP1 using randomly subsampled dataset (maximum 50 strains per variant; 474 VP1 sequences detected from 1995 to 2016). Data from the P domain is included here (amino acids 216-540). Residues were grouped into conserved sites (Shannon entropy value ≤ 0.3), sites mapping on newly defined variable motifs/antigenic sites (A-E, G, and H), and variable sites that mapped outside of the motifs/antigenic sites as in Fig. 2. (b) Based on structural analyses, four residues (250, 255, 300, and 329) were mapped on the surface of the VP1 protein outside of the motifs/antigenic sites. (c) Minor variation of the sequence patterns of those four sites suggested subtle impact of these residues on the variant emergence.

**Supplementary Figure 6: Mutagenesis analyses for antigenic site C mapping.** Differences among a Farmington Hills strain (MD2004-3 [DQ658413]), and a Sydney strain (RockvilleD1 [KY424328]) are shown for those residues from antigenic site C. Immunoassay performed with previously uncharacterized mAbs (B11 and B12) raised against the Farmington Hills strain (MD2004-3). Data from one mAb (B12) is shown in the context of different mutant VLPs from the Farmington Hills strain (MD2004-3). Mutation at residue 340 does not affect binding while progressive reduction of binding is detected when multiple substitutions are introduced in the VLPs.

**Supplementary Figure 7: Intra-variant mutational patterns of all the variable motifs/antigenic sites (A-E, G, and H) with legend**. (a) Amino acid pairwise differences among viruses from each of the GII.4 variants as compared to the earliest strain from each given variant. (b) Amino acids from each new and expanded motifs/antigenic sites were tabulated for each variant from GII.4 norovirus. Colors of the bars for the profiling graphs correspond to the predominant sequence pattern presented in that motif/antigenic site for each GII.4 variant. Patterns of the bars represent minor variations of sequences in the motifs/antigenic sites. The mutational patterns of each motif/antigenic site were listed in the legend of each bar graph.

**Supplementary Figure 8: Diversifying pressure on GII.4 intra-variant evolution.** Maximum likelihood trees of major GII.4 variants showing the diversifying pressure during the evolution. The branches under diversifying selection were explored using Mixed Effect Model of Evolution method (empirical Bayes factor > 100 and *p* < 0.05) and indicated in red.

**Supplementary Figure 9: Sequence pattern of NERK motif**. GII.4 variant distribution was plotted as described in Fig. 3. Amino acids from NERK motif was tabulated using 1572 sequences of GII.4 norovirus that circulated from 1995 to 2016. Colors of the bars for the graph correspond to the predominant sequence pattern presented in the NERK motif for each GII.4 variant.

**Supplementary Table 1: Dataset used in this study.**

